# Perigestational Opioid Exposure Alters Alcohol-Driven Reward Behaviors in Adolescent Rats

**DOI:** 10.1101/2023.11.14.567041

**Authors:** Christopher T. Searles, Hannah J. Harder, Meghan E. Vogt, Anne Z. Murphy

## Abstract

Every fifteen minutes, a baby is born in the U.S. experiencing neonatal opioid withdrawal syndrome (NOWS). Since 2004, the rate of NOWS has increased 7-fold. Clinical studies have established intrauterine exposure to drugs of abuse as a risk factor for adverse health outcomes in adult life, including the propensity for future illicit drug use. Despite extensive knowledge about common mechanisms of action in the neural circuitry that drives opioid and alcohol reward, there is little data on the risks that those born with NOWS face regarding alcohol use later in life. Here, we investigate the impact of perigestational opioid exposure (POE) on the mesolimbic reward system of male and female Sprague Dawley rats at postnatal and adolescent ages. Our laboratory has developed a clinically relevant model for morphine exposure spanning pre-conception to the first week of life. Using this model, we found that POE increased alcohol consumption in female rats under noncontingent conditions, and inversely, reduced alcohol consumption in both male and female rats during operant conditioning sessions. Operant responding was also reduced for sucrose, suggesting that the impact of POE on reward-seeking behaviors is not limited to drugs of abuse. Expression of µ-opioid receptors was also significantly altered in the nucleus accumbens and medial habenula, regions previously shown to play a significant role in reward/aversion circuitry.

**Significance Statement:** Early life exposure to opioids is known to alter future drug behavior in rats. In the present study, female rats exposed to morphine via their mothers throughout and after pregnancy exhibited increased alcohol consumption when allowed to consume freely. During operant conditioning, however, male and female rats exposed to gestational morphine decreased consumption of alcohol as well as sucrose. We also observed that gestational morphine exposure altered µ-opioid receptor expression in reward-related brain regions. Our study provides the first evidence of changes in alcohol-directed reward behavior in a gestational opioid exposure rat model.

## Introduction

Since 2004, the incidence of neonatal opioid withdrawal syndrome (NOWS) has increased almost 7-fold in the United States, most recently reaching 30,000 diagnoses per year (Patrick et al., 2015; Patrick et al., 2019; Winkleman et al., 2018). Pregnant women are prescribed opioids at alarmingly high rates (approximately 9.5%-41.6% of women in the U.S. alone), and it is currently estimated that a baby is born every 15 minutes with symptoms of opioid withdrawal (Desai et al., 2014; Winkelman et al., 2018). Rates of opioid abuse have also risen in adolescents, and among those seeking treatment, family history of opioid abuse has been identified as a primary contributing factor (Palamar et al., 2018; Russell et al., 2015). Indeed, familial drug use positively correlates with use in future generations in both humans and rodents (Cadoret et al., 1986; Vassoler et al., 2016). Gestational opioid exposure in humans is associated with several negative developmental outcomes that may further contribute to drug use, including cognitive deficits, reduced socialization, and increased anxiety (Azuine et al., 2019; Larson et al., 2019). This raises concern for drugs of abuse that are easily accessible, such as alcohol. Indeed, by age 18, nearly 60% of teens will have had at least one drink, with the majority having participated in binge-level consumption (SAMSHA, 2016).

Although chronic drug use has been shown to alter drug reward salience, limited research has explored the effects of POE on reward. Increased conditioned place preference for morphine was observed in adult male rats exposed to morphine during gestation (Torabi et al., 2017; Wu et al., 2009). Adult male rats exposed to morphine for one week of gestation also exhibited increased self-administration of morphine compared to controls (Torabi et al., 2017). Similar changes have been reported for POE and future use of methamphetamines and cocaine, suggesting that changes in sensitivity to drug reward span multiple drug classes (Ramsey et al., 1993; Shen et al., 2016; Vathy et al., 2003). It is unclear if this change in reward sensitivity is limited to drugs of abuse or extends to natural rewards (Gagin et al., 1996; Vathy et al., 2007). Although together these studies suggest that gestational exposure to morphine alters the reward value for drugs of abuse, many use a truncated exposure period, typically in the last week of pregnancy; thus, significantly limiting their translatability (Grecco & Atwood, 2020; Harder & Murphy, 2019). Our studies address this limitation by employing a clinically relevant model in which morphine is administered prior to and throughout gestation, extending through the first post-partum week. By initiating morphine exposure prior to mating, we mimic the typical user profile that develops into an opioid use disorder before pregnancy. Extending morphine exposure through seven days postnatally further replicates the use of supplemental opioid therapy that protects newborns from withdrawal and promotes the continuation of breastfeeding despite ongoing drug use (Abdel-Latif et al., 2006; Bogen & Whalen, 2019; Welle-Strand et al., 2013).

Using our clinically relevant POE model, we investigated the impact of perigestational morphine exposure on alcohol consumption and reward behaviors during adolescence and early adulthood. Our data reveal that perigestational morphine leads to effort-dependent changes in alcohol and sucrose consumption. These findings confirm that opioid use during pregnancy alters offspring’s risk of drug abuse across drug classes.

## Materials and Methods

### Subjects

Female Sprague Dawley rats (approximately two months of age) were used to generate male and female offspring perigestationally exposed to morphine or sterile saline for controls (Charles River Laboratories, Boston, MA). Rats were co-housed in same-sex pairs or groups of three on a 12:12 hours light/dark cycle (lights on at 8:00 AM). All rats were housed in Optirat GenII individually ventilated cages (Animal Care Systems, Centennial, Colorado, USA) with corncob bedding. Food (Lab Diet 5001 or Lab Diet 5015 for breeding pairs, St. Louis, MO, USA) and water were provided ad libitum throughout the experiment, except during operant procedures. All studies were approved by the Institutional Animal Care and Use Committee at Georgia State University and performed in compliance with the National Institutes of Health Guide for the Care and Use of Laboratory Animals. All efforts were made to reduce the number of rats used in these studies and minimize pain and suffering.

### Perigestational opioid exposure design

Female Sprague Dawley rats were implanted with iPrecio SMP-200 microinfusion minipumps at postnatal day 60 (P60). After one week of recovery from surgery, morphine administration began. Initially, rats were administered 10 mg/kg across three doses per day, with doses increasing weekly by 2 mg/kg/day until 16 mg/kg/day is reached. One week after morphine initiation, females were paired with sexually-experienced males for two weeks to induce pregnancy. Morphine exposure to the dams continued throughout gestation. At E18, pumps switched to twice-a-day dosing, as initial pilots using three-times-a-day dosing led to increased pup mortality early in life, possibly due to respiratory depression (data not shown). Dams continued to receive morphine after parturition, such that pups received morphine through their mother’s milk. Beginning at P5, the morphine dose was decreased by 2 mg/kg daily until P7, when the dose reaches 0 mg/kg. This protocol closely mirrors the clinical profile of opioid use disorder and neonatal opioid withdrawal syndrome. An essential part of this model is the inclusion of morphine dosing before E15, the approximate date of μ-opioid receptor development in the fetal brain (Coyle and Pert, 1976), as there are potential trophic effects of opioids early in development (Kuhn et al., 1992). Pumps filled with sterile saline were used to generate control rats to account for the stress of surgery and pump refilling.

### Alcohol exposure

Male and female rats exposed to saline or morphine during gestation were weaned into reverse light conditions (12:12 hours dark/light; lights on at 8:00 PM) at P21 and aged to adolescence undisturbed before beginning daily alcohol exposure at P30.

Alcohol consumption was assessed using self-administration of ethanol gelatin (10% ethanol, 8% Polycose, and 2.5% gelatin in tap water). Consumption of gelatin generates blood and brain ethanol levels in Sprague-Dawley rats that are comparable to other paradigms involving liquid delivery (Nurmi et al., 1999; Peris et al., 2006). The protocol also avoids the necessity for alcohol-preferring rats that may exhibit different baseline expression of µ-opioid receptor underlying increased consumption (McBride et al., 1998). Eighteen male (11 saline vehicle and 7 morphine) and sixteen female (9 saline vehicle and 7 morphine) rats began daily alcohol exposure with conditioned place preference (CPP) conditioning and were used for analysis. Two to five rats per group were excluded from CPP analysis but underwent all testing. Twenty-eight male (15 saline vehicle and 13 morphine) and twenty-five female (11 saline vehicle and 14 morphine rats) began daily alcohol exposure with noncontingent sessions. No differences were found in alcohol consumption between groups that underwent CPP conditioning and those that did not, so data was pooled by matching age of experience. For this reason, sample size at Days 2 and 4 of the noncontingent sessions is decreased.

### Conditioned place preference

Two types of bedding (both distinct from the rats’ typical home-cage bedding) were used as environmental indicators. CPP testing took place under red-light in a two-compartment arena separated by an open door; one side contained white paper square bedding, whereas the other side had pelleted paper bedding (Teklad, Envigo, Indianapolis, IN). Rats were habituated to the testing room for approximately one hour before testing. On P30, rats were recorded exploring the arena freely for 20 minutes as a pretest. Side preference was determined (>50% time spent) and used to generate pseudorandom unbiased and counterbalanced assignments. Half of the animals were assigned alcohol exposure with bedding from their preferred side and half with bedding of their non-preferred side. Rats exhibiting >70% preference for one side during the pre-conditioning test were excluded from CPP analysis, but not removed from further experiments. Conditioning sessions took place in a home cage-like setting with paired bedding. During alcohol-paired sessions, one of two ad libitum water bottles was replaced with a jar of approx. 20g of ethanol gelatin. Sessions without alcohol contained two water bottles and the other bedding type. Conditioning began on Day 1 with 24-hour exposure to one bedding type. Following this were a second 24-hour conditioning session, two days with one 6-hour session, two days with two 3-hour sessions, and five days with two 1-hour sessions. Three-hour and one-hour conditioning sessions took place twice daily, separated by 3 hours. Order of alcohol/no alcohol sessions was randomly assigned and then alternated each day (Figure 2a). Days with two sessions alternated morning/afternoon:alcohol/no alcohol to balance for alcohol exposure time of day. Rats were single-housed only during CPP conditioning and were otherwise group-housed. The post-conditioning test session was conducted on the 12th day. During the test, rats were placed in the center doorway randomly facing an arena side/bedding type. Rats were recorded exploring the arena freely for 20 minutes. Preference scores were calculated by subtracting the time spent on the non-alcohol-paired side from the time spent on the alcohol-paired side. Percent change in preference score between pre- and post-conditioning tests was also calculated.

**Figure 1.**
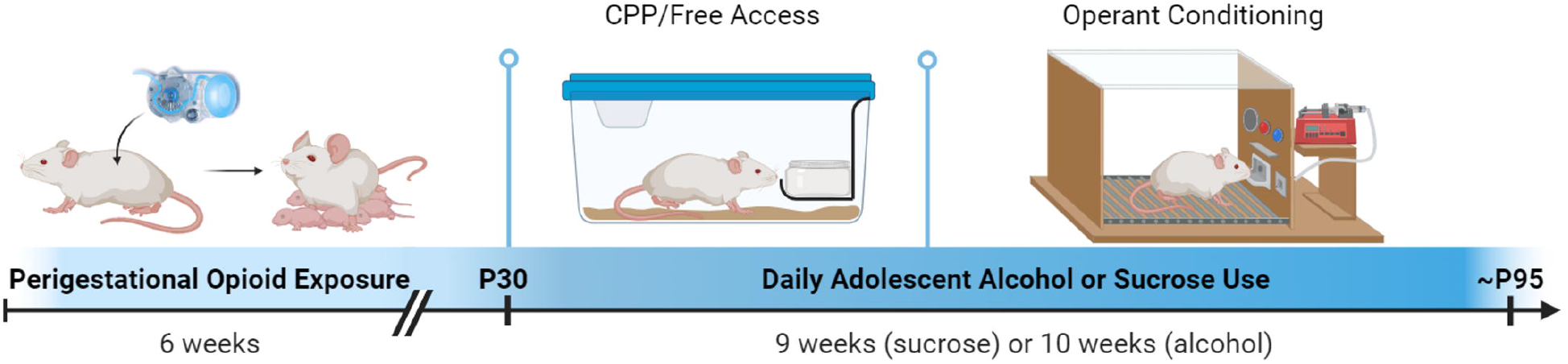
Sprague Dawley rats are exposed to morphine or saline via their mothers throughout three weeks of gestation and one week following birth via their mother’s milk. Morphine or saline exposure for mothers begins prior to breeding, resulting in approximately one week of saline (all rats) and five weeks of morphine (POE). Beginning at P30, drug-exposed experimental rats begin alcohol or sucrose self-administration sessions that last for several weeks. Created with Biorender.

**Figure 2.**
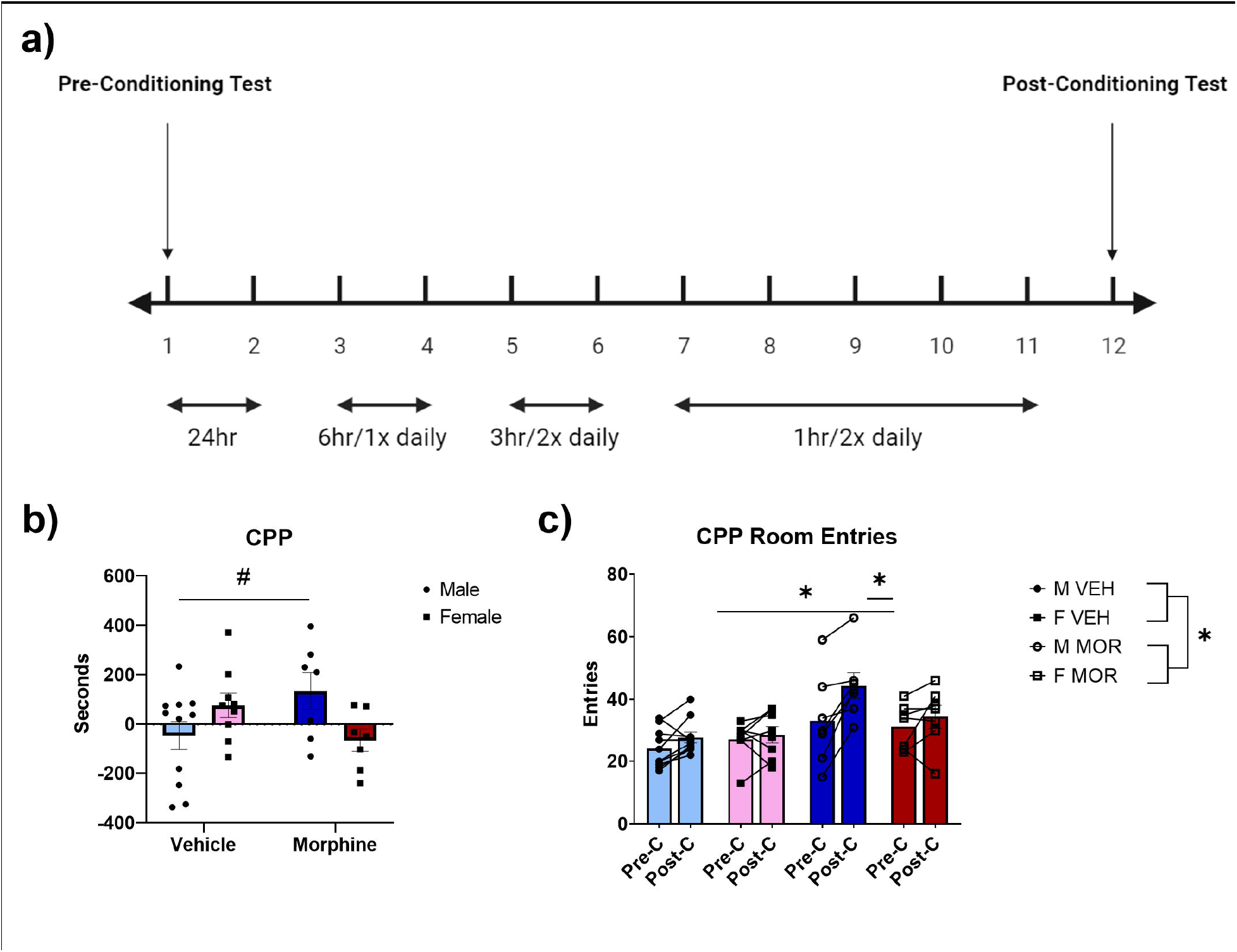
Conditioned Place Preference for alcohol begins at P30 and takes place across twelve days (**a**). POE male rats showed a trending increase in seconds spent post-conditioning on the alcohol-paired side of the arena compared to vehicle males, indicating an increased preference for alcohol (**b**). POE male rats also increased their exploratory behavior during CPP testing compared to vehicle males, as demonstrated by an increase in room entries (**c**). Overall POE rats showed increased exploratory behavior. #Trending difference between POE males vs vehicle males; p=0.06. *Significant difference between POE male pre-conditioning vs post-conditioning, POE male post-conditioning vs vehicle males post-conditioning, or POE rats vs vehicle control rats; p < 0.05. Graphs indicate mean ± SEM.

### Noncontingent alcohol self-administration

Similar to CPP, rats were isolated in a home cage-like setting with a jar filled with ethanol gelatin for two consecutive 24-hour sessions. Free access to ethanol gelatin was gradually reduced across the next 4 days to 1-hour daily sessions that continued for 10 total days (Figure 3a). Ethanol consumption was measured in g/kg/hr. Given that CPP and non-CPP home-cage access protocols were identical in length and within-group data did not statistically differ or differentially impact future behaviors, these data were combined.

**Figure 3.**
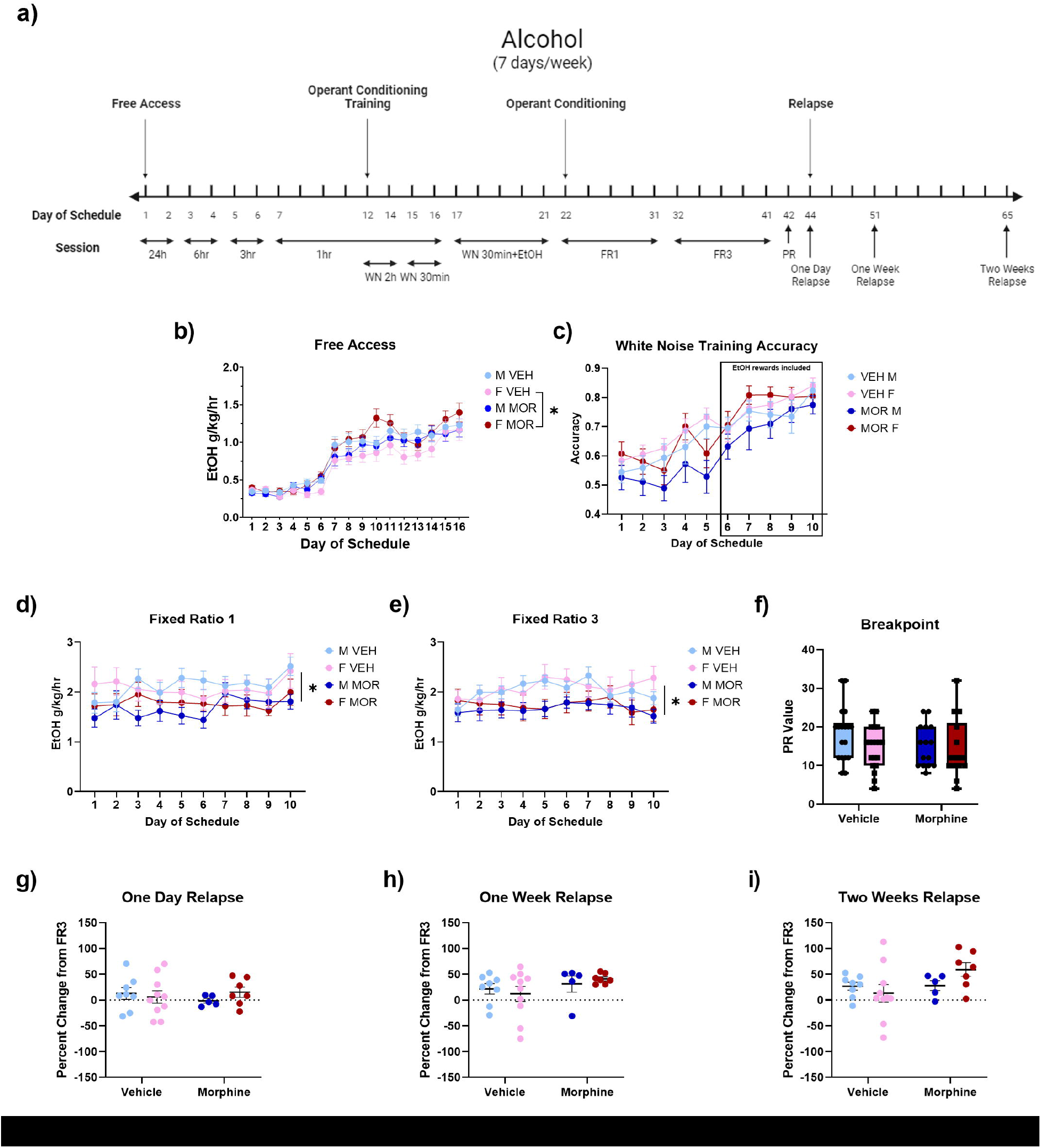
Our alcohol consumption schedule takes place across 65 days (**a**). POE female rats consumed more g/kg/hr of alcohol than vehicle females in a noncontingent, home-cage-like setting (**b**). Groups did not differ in active lever press (AP) accuracy during training; accuracy is defined by (AP / (AP during reward receipt + inactive presses)) (**c**). POE rats consumed significantly less alcohol during operant conditioning sessions at both FR1 and FR3 schedules (**d-e**). Groups did not differ in breakpoint for alcohol as determined by progressive ratio schedule (**f**). Groups did not significantly differ in relapse potential; values indicate individual % change in alcohol consumed from 10-day average of FR3 (**g-i**). Increase in percent change suggests relapsed alcohol consumption. *Significant difference between POE females vs vehicle females, or POE rats vs vehicle control rats; p<0.05. Graphs indicate mean ± SEM.

### Operant procedures

During the last five days of noncontingent regimen, rats were trained to lever press in operant conditioning chambers using light cues and negative reinforcement (80db white noise off for five seconds). A light cue indicated successful pressing of the “active” (versus inactive) lever. After five days of negative reinforcement, gelatin rewards were paired with the reinforcement cues. All rats underwent ten days of 30-minute sessions where an active lever press earned an approximately 0.2-gram ethanol gelatin reward (Fixed Ratio 1 or FR1), followed by ten days of FR3 in which three presses are required for rewards. On the final day, motivational breakpoints were determined using a progressive ratio (PR) in which reward requirements increase as each is achieved. Record of the last reward achieved indicates motivation to obtain (or “wanting”) alcohol (Berridge, 1996). Throughout operant conditioning sessions, lever press data from both active and inactive levers was collected using time signatures to monitor activity levels, total consumption, consumption across time, and lever differentiation. Within the chambers, infrared beams at the front and back of the chambers monitored animal activity during sessions.

### Relapse testing

Forced abstinence is a well-established behavior test used to measure the rewarding properties of alcohol after daily use is suspended (Lê & Shaham, 2002). Following breakpoint determination, rats were returned to their home cage for one day, one week, and two weeks of abstinence. Following abstinence, gelatin consumption was measured in one FR3 operant session.

### Sucrose Administration

A separate cohort of rats was tested for reward behavior towards 20% sucrose. Sucrose was mized with 2.5% gelatin in tap water and administered in a home cage-like setting using a shortened schedule described for ethanol gelatin (Figure 4a). Sucrose was administered in tap water without gelatin during operant conditioning with a similar schedule to ethanol gelatin (Figure 4a). Sucrose sessions took place only five days a week. Follow-up PR and relapse testing also took place as described for ethanol gelatin without a one-day relapse test given the regular two-day breaks in the sucrose schedule.

**Figure 4.**
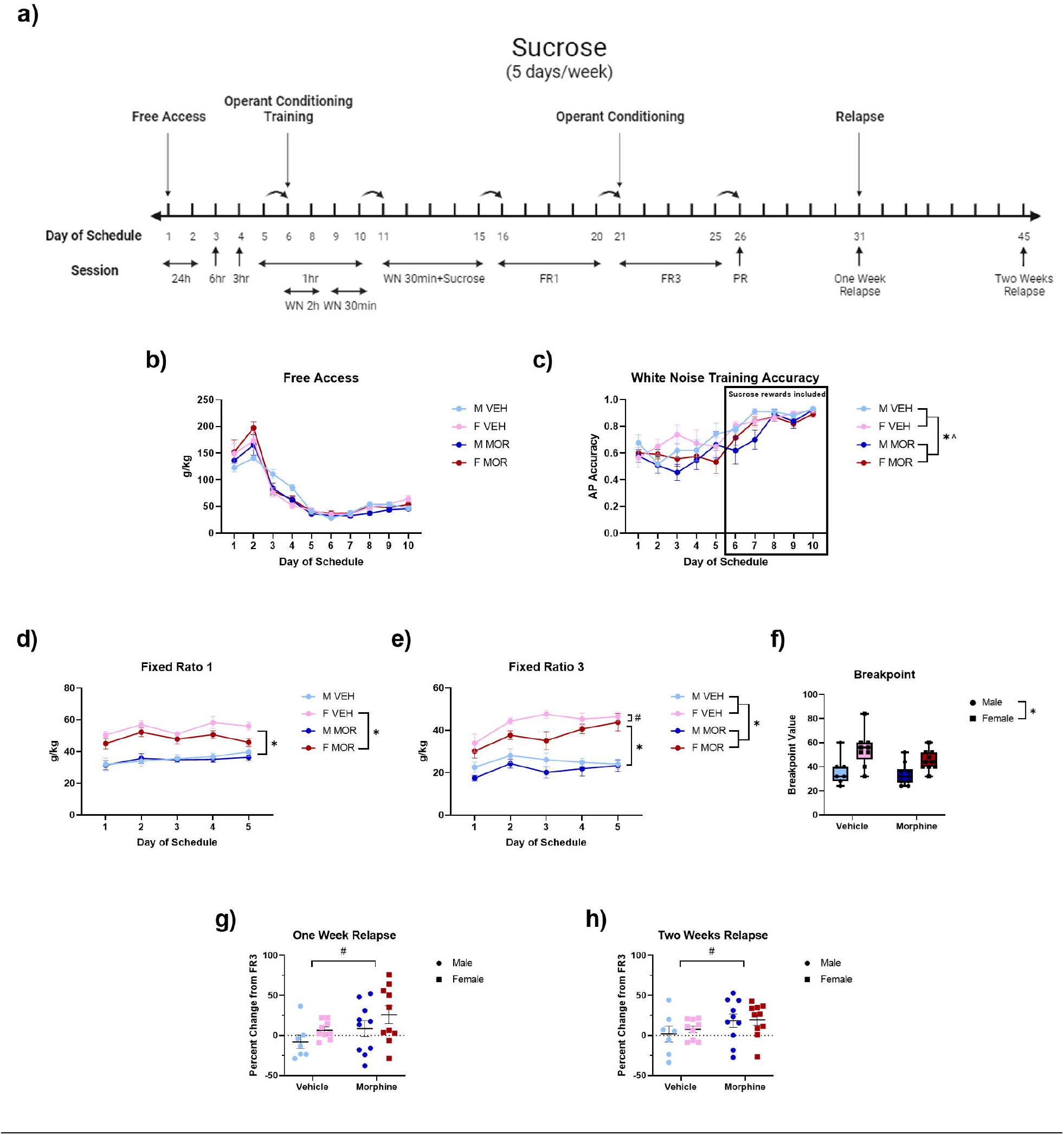
Our sucrose consumption schedule takes place across 45 days, arrows indicate 2-day breaks in behavioral testing (**a**). Groups did not differ in sucrose consumption in a noncontingent, home-cage-like setting (**b**). Vehicle rats were more accurate than morphine rats in their active lever press (AP) accuracy during training; accuracy is defined by (AP / (AP during reward receipt + inactive presses)) (**c**). This is also true when considering only the last five days of training where sucrose rewards were included. POE female rats consumed significantly less sucrose than control females during operant conditioning session at FR1 (**d**) and showed a trending decrease at FR3 (**e**). During FR1 and FR3, female rats overall consumed more sucrose than male rats. Vehicle rats overall consumed more sucrose during FR3 than POE rats. Female rats exhibited higher breakpoints for sucrose than male rats as determined by progressive ratio schedule (**f**). POE rats showed a trending increase in relapse potential compared to vehicles after one week and two weeks forced abstinence; values indicate individual % change in sucrose consumed from 10-day average of FR3 (**g-h**). Increase in percent change suggests relapsed drinking. *Significant difference between POE rats vs vehicle control rats, female rats vs male rats, or POE females vs vehicle females; p<0.05. ^Significant difference between POE rats vs vehicle rats across Days 5-10; p<0.05. #Trending difference between POE females vs vehicle females or POE rats vs vehicle rats; p<0.08. Graphs indicate mean ± SEM.

### Immunohistochemistry

Randomly selected littermates of rats used for reward behavior were sacrificed at postnatal ages 7, 14, and 30 for assessment of µ-opioid receptor (MOR) expression. Brain tissue was collected via rapid decapitation and drop-fixed in 4% paraformaldehyde for 24 hours, followed by storage in 30% sucrose at 4°C until sectioning. Fixed tissue was sectioned coronally at 40-µm with a Leica SM2010R microtome and stored at −20°C in cryoprotectant-antifreeze solution (Watson et al., 1986). Forebrain and midbrain sections were processed for MOR immunoreactivity using standard techniques previously described (Harder et al., 2023). Free-floating sections were rinsed in potassium phosphate-buffered saline (KPBS), incubated in 3% hydrogen peroxide for 30 minutes, and rinsed again in KPBS at room temperature. Tissue was then incubated in rabbit anti-MOR primary antibody (Alomone Labs, Jerusalem, Israel; 1:7,500 for P7 and P14; 1:2,000 for P30) at room temperature for 1 hour followed by 48 hours at 4°C. Primary antibody was rinsed using KPBS and incubated in biotinylated donkey anti-rabbit IgG (Jackson Immunoresearch, West Grove, PA, USA; 1:600) for 1 hour. After tissue was rinsed with KPBS and sodium acetate (0.175M; pH 6.5), immunoreactivity was visualized using nickel sulfate 3,3′-diaminobenzidine solution (2 mg/10 ml) and 0.08% hydrogen peroxide in sodium acetate buffer. After 20 minutes, tissue was rinsed in sodium acetate and KPBS before being mounted on gelatin-subbed slides, air-dried, dehydrated with increasing concentrations of ethanol, cleared with xylene, and cover-slipped.

For immunohistochemistry data, 16-bit grayscale images were captured using a QImaging Retiga EXi CCD camera and IPLab Image Analysis Software (BD Biosciences). The nucleus accumbens (NAc), medial habenula (MHb), and ventral tegmental area (VTA) were bilaterally sampled at 10× magnification 2-3 times per subject (4-6 values) and the average grayscale pixel value within each region was recorded.

### Statistical analysis

Significant effects of sex and morphine treatment were assessed using two- or three-way mixed models or repeated measures mixed models with an alpha level of p<0.05. Given that ANOVA testing cannot handle missing values, mixed models using Greenhouse-Geisser were implemented where outlier correction was necessary. Outliers were discovered using GraphPad Prism’s ROUTS method. Given that findings of previous POE and drug abuse literature are often sex-specific, *a priori* specified tests comparing male and female groups independently were performed using t-test or ANOVA to determine significant mean differences. Other post hoc testing was performed using Tukey’s or Sidak’s. Experiments involving repeated measures were not tested for daily individual group differences. Individual litter differences were not found; therefore, group data are comprised of multiple litters. GraphPad Prism 9.1.0 was used for all statistical analysis.

## Results

### Conditioned place preference

Rats exposed to morphine (POE) or saline (vehicle control) exposure were initially tested under CPP conditions for alcohol preference. POE male rats trended towards increased time spent in the alcohol-paired bedding area after conditioning compared to vehicle control males (F_interaction_(1,30)=7.50, p=0.01; t(16)=1.947, p=0.06; Figure 2b). POE male rats also exhibited increased postconditioning room entries, or movement between the two arena areas, compared to the preconditioning test (t(6)=3.994, p<0.01; Figure 2c). Alcohol conditioning did not have this effect on other groups. POE males exhibited significantly more room entries compared to control males during post-conditioning testing (Tukey’s HSD, p<0.01, 95% CI=[- 29.85 to −3.124]). Overall, POE rats exhibited more room entries than control rats (F_treatment_(1,28)=9.66, p<0.01). Groups did not significantly differ in consumption of alcohol per hour during conditioning, though POE females trended towards increased consumption compared to control females (F_treatment_(1,44)=3.9, p=0.0548; Supplemental Figure 2a).

### Noncontingent Alcohol Self-Administration

Males and females did not differ across treatment in body weight throughout daily alcohol self-administration sessions (F_males_(1,51)=3.25, p=0.08, F_females_(1,46)=0.10, p=0.75; Supplemental Figure 1). Rats of all groups were introduced to alcohol in “free access” home-cage-like isolation where self-administration was allowed for initially prolonged and then only brief periods of time each day. Across these sessions, POE female rats consumed significantly more grams of pure alcohol per hour compared to control females (F_sex*treatment interaction_(1,97)=4.52, p=0.036; Figure 3b). ANOVA of only female data supports this result (F_treatment_(1,46)=4.194, p=0.04).

### Contingent Alcohol Self-Administration

Across the last five days of noncontingent consumption, rats underwent alcohol reward-free operant lever press training using white noise reinforcement and light cues. Training continued for an additional five days with alcohol rewards included with conditioned cues (Figure 3a). Across these sessions, groups did not differ in lever pressing behavior (Supplemental Figure 2b). Groups were also similar in overall lever press accuracy, though may have exhibited differences on individual days (F_day*treatment interaction_(1,63)=2.072, p=0.04; Figure 3c). Female rats overall were more accurate than male rats in their press accuracy, regardless of treatment (F_sex_(1,63)=5.050, p=0.028). After training, earned gelatin rewards were accompanied only by light cues. Across ten days of FR1 operant conditioning, POE rats consumed significantly less alcohol compared to vehicle controls (F_treatment_(1,63) = 6.424, p=0.01; Figure 3d). This reduced alcohol consumption continued through ten subsequent days of FR3 (F_treatment_(1,63)=4.805, p=0.03; Figure 3e). *A priori* ANOVA determined that these differences were driven primarily by male groups (F_maleFR1treatment_(1,31)=7.595, p<0.01, F_maleFR3treatment_(1,31)=4.697, p=0.03). Despite differences in consumption, groups did not differ in number of rewards achieved across the duration of the average session (Supplemental Figures 2g-h). Groups did not differ in maximal effort, as measured by breakpoint, when tested under progressive ratio conditions (Figure 3f). Relapse was determined by changes in consumption behavior after successive forced abstinence periods of one day, one week, and two weeks. Groups did not differ in relapse potential, or consumption after abstinence compared to individual average FR3 consumption (Figure 3g-i). However, unlike other groups, all of our POE female rats exhibited relapsed drinking behavior at one week’s and two weeks’ testing.

Data gathered from operant conditioning sessions also provided information on task focus. Although not consistent across sessions, POE females earned their first FR3 reward faster than vehicle females (F_day*treatment interaction_(9,540)=2.052, p=0.032; Supplemental Figure 2c). Males were overall faster to earn rewards than females (F_sex_(1,63)=11.79, p<0.01). Males were also more accurate in their lever pressing compared to females (F_sex_(1,63)=6.865, p=0.01, Supplemental Figure 2d). Accuracy is represented as the percentage of all lever presses that elicited a reward; errant pressing includes those on the active lever that did not work towards a reward. Sex differences also appeared in relapse testing, in that males were faster to earn their first reward than females (F_sex_(1,63)=6.865, p<0.01; Supplemental Figure 2e). After one day of forced abstinence, POE rats were faster to their rewards than vehicle rats (F_day*treatment interaction_(2,54)=4.043, p=0.02). Sex differences in lever press accuracy did not extend to relapse testing (Supplemental Figure 2f).

### Sucrose Self-Administration

Sucrose self-administration was used to determine if changes found in alcohol-driven behaviors due to POE extend to natural reward. Groups did not differ in their noncontingent consumption of 20% sucrose (Figure 4b). During operant conditioning training using white noise, vehicle rats were more accurate in their active lever pressing than POE rats (F_treatment_(1,32)=5.717, p=0.023; Figure 4c). This significant difference is maintained when considering only the last five days of training where sucrose rewards were included (F_treatment_(1,32)=4.260, p=0.047). Across FR1 sessions, female rats of both treatments consumed significantly more sucrose than males (F_sex_(1,32)=63.51, p<0.01, Figure 4d). POE female rats also consumed significantly less sucrose than vehicle females according to *a priori* ANOVA (F_treatment_(1,32)=63.51, p=0.046). During FR3 sessions, there was both a treatment and sex difference, in that female rats continue to consume more sucrose than males overall, while vehicle groups consumed more than morphine groups (F_sex_(1,32)=56.24, p<0.01, F_treatment_(1,32)=4.489, p=0.042; Figure 4e). *A priori* two-way ANOVA, however, revealed that treatment differences were not conserved when considering each sex independently (F_females_(1,17)=3.567, p=0.076, F_males_(1,15)=1.271, p=0.277). During FR1 and FR3, all groups exhibit similar reduction in sucrose rewards achieved across the average session (Supplemental Figure 3f-g). Sucrose breakpoint as measured by progressive ratio testing was higher in females compared to males but not altered by POE treatment (F_sex_(1,32)=16.37, p<0.01, F_treatment_(1,32)=2.872,p=0.099; Figure 4f). This difference is also reflected in total active presses (F_sex_(1,32)=12.88, p<0.01, F_treatment_(1,32)=3.043, p=0.09; not shown). After one week forced abstinence, POE rats trended towards increased relapse potential compared to vehicle rats, demonstrating higher increases in sucrose consumption compared to their average FR3 consumption (F_treatment,_ (1,32)=3.793,p=0.0603; Figure 4h). This was also true after a subsequent two-week forced abstinence period (F_treatment_(1,32)=3.522,p=0.0697; Figure 4i). During FR3, press accuracy towards sucrose was decreased in females but not different between treatment groups (F_sex_(1,32)=11.40, p<0.01, F_treatment_(1,32)=3.109, p=0.087; Supplemental Figure 3c). POE treatment did not alter first reward latency of sucrose during FR3 (F_treatment_(1,32)=0.02408, p=0.88; Supplemental Figure 3b). During relapse, males of both groups were more accurate than females in their lever pressing, but did not differ in first reward latency (F_sex_(1,32)=5.325, p=0.0276, Supplemental Figure 3d-e).

### µ-Opioid Receptor Expression

Group and sex differences in expression of µ-opioid receptor (MOR) protein was analyzed using immunohistochemistry. Key brain regions involved in reward and three early life time points were considered. At P7, there were no group differences in MOR expression in the nucleus accumbens (NAc), medial habenula (MHb), or ventral tegmental area (VTA) (Figure 5a, d, g). At P14, POE females expressed increased NAc MOR compared to vehicle females and POE males (F_sex*treatment interaction_(1,19)=13.42, p<0.01; Figure 5b). No group differences in MHb or VTA MOR expression were found at P14 (Figure 5e, h). At P30, POE groups expressed significantly less MHb MOR than vehicle groups (F_treatment_(1,19)=4.848, p=0.0402; Figure 5f). No group differences in NAc or VTA MOR expression were found at P30 (Figure 5c, i).

**Figure 5.**
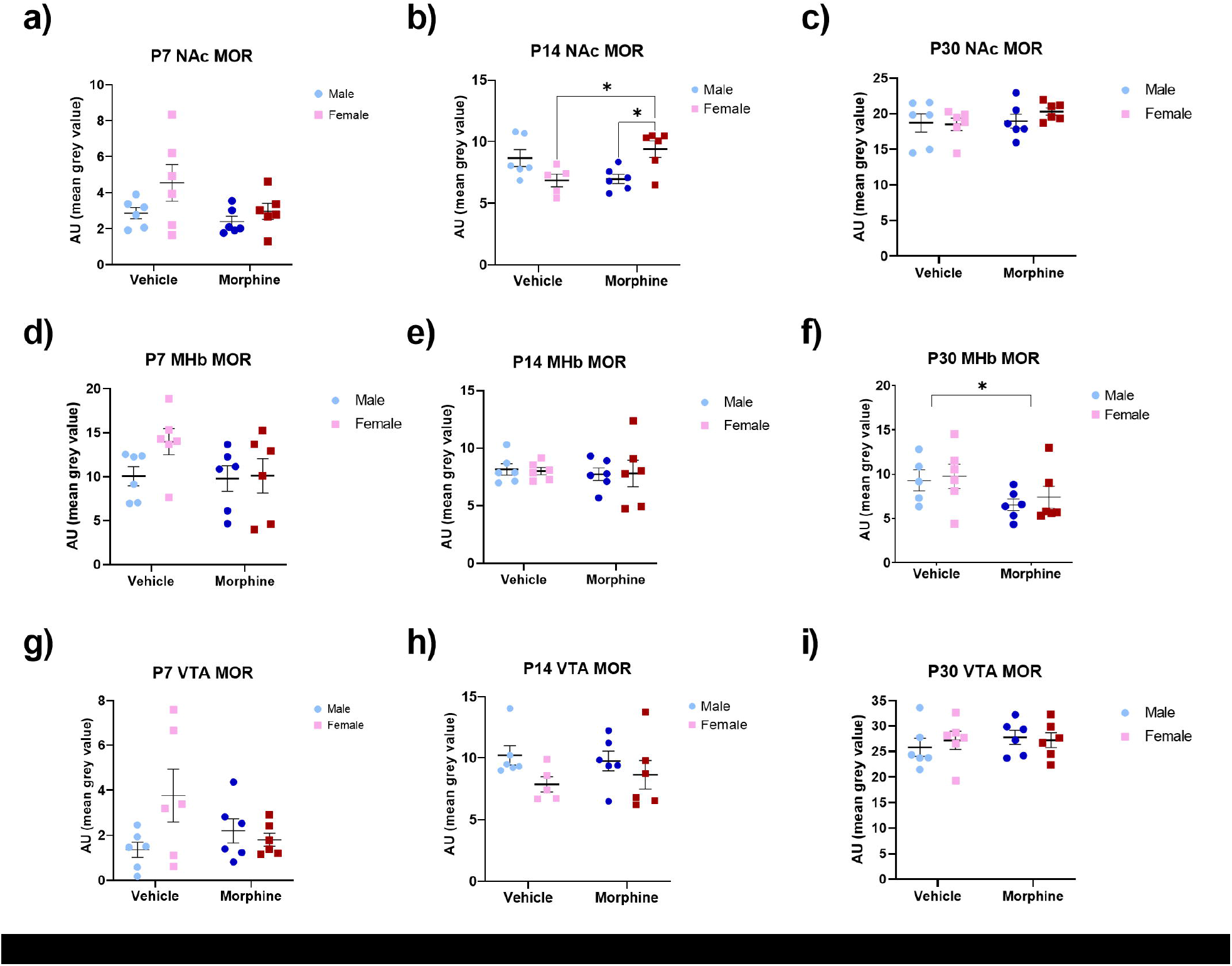
µ-Opioid receptor expression in nucleus accumbens (NAc), medial habenula (MHb), and ventral tegmental area (VTA) during early life. NAc MOR expression is elevated in POE female rats compared to vehicle female rats at P14, but no other groups differences were across P7, P14, or P30 (**a-c**). MHb MOR expression is significantly lower in POE rats compared to vehicle rats at P30, but no differences were found at P7 or P14 (**d-f**). No group differences were found in VTA MOR across P7, P14, or P30 (**g-i**). *Significant difference between POE female rats vs vehicle female rats or POE rats and vehicle rats; p<0.05. Graphs indicate mean± SEM.

## Discussion

Our results show that perigestational morphine exposure alters multiple alcohol-driven behaviors in both male and female rats. This is the first time that perigestational opioid exposure has been shown to impact alcohol consumption in the rat model. Additionally, alcohol-related differences in POE rats were in some cases sex-dependent. POE altered CPP behaviors including preference for alcohol-paired cues and alcohol-seeking locomotion in males. Increased alcohol consumption was found amongst POE female rats compared to vehicle female rats during noncontingent, home-cage-like exposure sessions. This increase did not extend to operant conditioning sessions where additional effort was required to earn gel rewards. Surprisingly, POE rats significantly reduced their alcohol consumption across FR1 and FR3 compared to control rats. This was also true for our natural reward choice, sucrose, where POE rats consumed less than vehicle controls during FR3. POE rats also displayed elevated relapse behaviors towards sucrose.

Our data show that POE-induced changes in alcohol consumption are not consistent across drinking models, contrary to previous, preclinical studies that have shown enhanced sensitivity and motivation for drugs of abuse after early life exposure to opioids (Wu et al., 2009; Ramsey et al., 1993; Shen et al, 2016; Chiang et al., 2014; Vassoler et al., 2016). Outcomes of previous studies may have been exaggerated by cross-fostering or early life withdrawal precipitation using MOR antagonists, such as naloxone (Gagin et al., 1996; Torabi et al., 2017). These practices produce a more aggressive withdrawal profile compared to the recommended clinical treatment of supplemental opioid agonists (Abdel-Latif et al., 2006; Bogen & Whalen, 2019; Kocherlakota, 2014; Welle-Strand et al., 2013). The use of truncated opioid exposure periods, typically in the last week of pregnancy, significantly limits the translatability of some POE models (Grecco & Atwood, 2020). Our results may also be due to alcohol’s variable reward value in rats compared to other drugs of abuse, such as stimulants that are highly addictive. Rats are known to less readily consume alcohol compared to other drugs, normally necessitating the use of preferring “P” strains. Particularly, significant drinking behavior during adolescence in non-selectively bred rats is typically unlikely (McBride et al., 2014). Our Sprague Dawley rats, however, consumed at high enough rates (>1g/kg/hr) to be representative of adverse and binge alcohol drinking in humans as defined by U.S. Environmental Protection Agency guidelines and National Institute on Alcohol Abuse and Alcoholism reports respectively (>531mg/kg, Lachenmeier & Rehm, 2015; >1mg/kg, NIAAA). Consumption in excess of 1g/kg/hr leads to blood alcohol concentrations of at least 0.08%, or 80mg/dL (Peris et al., 2006).

Despite the self-administered nature of alcohol conditioning in our model, rats did not differ in amount consumed (Supplemental Figure 2a). Increased reward-context seeking during CPP suggests that POE may produce increased sensitivity to the rewarding effects of alcohol. Generating CPP for alcohol in rats is uncommon and requires additional experience or selective-breeding. Typically, exposure to alcohol produces aversion or no preference (Roger-Sánchez et al., 2012). Our control males exhibit this behavior (Figure 2b). For females, reward-context appears more important during operant conditioning where POE produced consistently elevated alcohol relapse. This relapse also had a trending increase as abstinence time increased, suggesting that females may be more susceptible to context-reinstatement behaviors when alcohol is present and when abstinence is prolonged. Negative affect, which is more prevalent in humans and rats exposed to perigestational opioids, is perpetuated by abstinence and can lead to relapse behavior (Bakhireva et al., 2019; Konijnenberg et al., 2016; Castro et al., 2022; Koob, 2020). Symptoms of negative affect produced through chronic stress, however, can lead to opposing reward-seeking changes. Chronic social isolation has been shown to increase two-bottle choice alcohol consumption while chronic social defeat and chronic restraint stress reduce operant responding for sucrose in rodents (Lee et. al, 2021; Bergamini et al., 2016; Xu et al., 2017). These inverse results were seen in our study in which POE rats consumed more alcohol compared to control rats during noncontingent or “free access” sessions but less during operant conditioning sessions. Importantly, decreased alcohol consumption during operant conditioning was not a result of impaired learning as shown by lever press accuracy during training (Figure 3c). Motivational disturbances can accompany both mesolimbic dopamine dysfunction and anhedonic phenotypes in rats (Canonica & Zalachoras, 2022). These physiological and emotional outcomes could be important underlying contributors to our results and require future exploration.

Several reward behavior changes found with respect to alcohol were also found when using sucrose, a natural reward. In previous literature where sucrose has been used for reward training only, consumption was unaffected by POE (Vathy et al., 2007). Longer protocols, however, present inconsistent differences in behavior. In Gagin et al. (1996) male and female rats given escalating doses of morphine from gestational (G) day 12-18 and cross-fostered after birth displayed increased preference for saccharine. Torabi et al. (2017) found that escalating, and subsequent descending, doses of morphine from G11 to birth and naloxone-precipitated withdrawal resulted in decreased sucrose preference and increased morphine preference in male and female rats. With our prolonged daily sucrose exposure protocol, we found no treatment differences until operant conditioning in the fourth week of the protocol. Here, sucrose consumption was decreased in POE rats, especially females. Moreover, POE rats demonstrated increased relapse for sucrose compared to controls. These findings suggest that changes in reward-seeking behavior is not isolated to drugs of abuse.

Both sucrose and alcohol consumption in rodents is shown to be impacted by manipulation of MOR (Hayward et al., 2006; Heyser et al., 1999; Stromberg et al., 1996; Henderson-Redmond & Czachowski, 2014; Morales et al., 2020). POE-induced changes in MOR populations across brain regions specific to reward and stress-reactivity are well-characterized. Here, we found that at MOR expression was increased in the nucleus accumbens (NAc) of POE female rats at P14 (Figure 5b). NAc MOR is critical to the rewarding properties of alcohol in rats and its pharmacological manipulation leads to changes in consumption behaviors (Barson et al., 2009; Hyytia et al., 2001). Our finding aligns with previous studies showing POE-induced increases in NAc MOR expression and binding (Handelmann & Quirion, 1983; Vassoler et al., 2016; Vathy et al., 2003). Additionally, at P30, we found that medial habenula (MHb) MOR expression was decreased in POE rats compared to controls. The MHb has the highest region-specific expression of MOR in the brain and plays a significant role in opioid reward/aversion circuitry and social interaction (Boulos et al., 2017; Gardon et al., 2014; Mechling et al., 2016; Allain et al., 2022). We have previously shown that our POE model significantly reduced social interaction in periadolescent rats (Harder et al., 2023). Boulos et al. (2020) found that manipulation of MHb MOR contributed to changes in naloxone aversion, but not food responding, when striatal and midbrain MOR expression was conserved. Given that our changes in MOR are not isolated to MHb, we cannot rule out this region’s potential role in reported changes in behavior. It is also important that we examine MOR in other brain regions reportedly altered by POE, such as the amygdala and hippocampus (Vathy et al., 2003; Schindler et al., 2004).

Our study is the first to report that POE induces changes in alcohol and sucrose self-administration and does so using a biologically relevant opioid exposure model. Though we did not specifically test for changes in emotional states, altered motivation for sucrose specifically may suggest symptoms of anhedonia amongst POE rats. Future experimentation is necessary for understanding if POE produces anhedonia that may underlie observed behavioral changes.

## Acknowledgements

This work was supported by the National Institutes of Health (1RO1DA041529). The funding source had no involvement in study design; data collection, analysis or interpretation; manuscript writing; or the decision to publish this article. Morphine sulfate was provided by the NIDA Drug Supply Program.

## References

Abdel-Latif M. E., Pinner J., Clews S., Cooke F., Lui K., Oei J. (2006). Effects of breast milk on the severity and outcome of neonatal abstinence syndrome among infants of drug-dependent mothers. Pediatrics, 117(6), e1163–e1169. 10.1542/peds.2005-1561

Allain, F., Carter, M., Dumas, S., Darcq, E., & Kieffer, B. L. (2022). The mu opioid receptor and the orphan receptor GPR151 contribute to social reward in the habenula. Scientific reports, 12(1), 20234. 10.1038/s41598-022-24395-z

Azuine R. E., Ji Y., Chang H. Y., Kim Y., Ji H., DiBari J., Hong X., Wang G., Singh G. K., Pearson C., Zuckerman B., Surkan P. J., Wang X. (2019). Prenatal Risk Factors and Perinatal and Postnatal Outcomes Associated With Maternal Opioid Exposure in an Urban, Low-Income, Multiethnic US Population. JAMA network open, 2(6), e196405. 10.1001/jamanetworkopen.2019.6405

Bakhireva L. N., Holbrook B. D., Shrestha S., Leyva Y., Ashley M., Cano S., Lowe J., Stephen J. M., Leeman L. (2019). Association between prenatal opioid exposure, neonatal opioid withdrawal syndrome, and neurodevelopmental and behavioral outcomes at 5–8 months of age. Early Human Development, 128, 69–76. 10.1016/J.EARLHUMDEV.2018.10.010

Barson, J. R., Carr, A. J., Soun, J. E., Sobhani, N. C., Leibowitz, S. F., & Hoebel, B. G. (2009). Opioids in the nucleus accumbens stimulate ethanol intake. Physiology & behavior, 98(4), 453–459. 10.1016/j.physbeh.2009.07.012

Bergamini, G., Cathomas, F., Auer, S., Sigrist, H., Seifritz, E., Patterson, M., Gabriel, C., & Pryce, C.R. (2016). Mouse psychosocial stress reduces motivation and cognitive function in operant reward tests: A model for reward pathology with effects of agomelatine. European neuropsychopharmacology, 26(9), 1448–1464. 10.1016/j.euroneuro.2016.06.009

Berridge K. C. (1996). Food reward: brain substrates of wanting and liking. Neuroscience and biobehavioral reviews, 20(1), 1–25. 10.1016/0149-7634(95)00033-b

Bogen, D. L., & Whalen, B. L. (2019). Breastmilk feeding for mothers and infants with opioid exposure: What is best?. Seminars in fetal & neonatal medicine, 24(2), 95–104. 10.1016/j.siny.2019.01.001

Boulos, L. J., Darcq, E., & Kieffer, B. L. (2017). Translating the Habenula-From Rodents to Humans. Biological psychiatry, 81(4), 296–305. 10.1016/j.biopsych.2016.06.003

Boulos, L. J., Ben Hamida, S., Bailly, J., Maitra, M., Ehrlich, A. T., Gavériaux-Ruff, C., Darcq, E., & Kieffer, B. L. (2020). Mu opioid receptors in the medial habenula contribute to naloxone aversion. Neuropsychopharmacology, 45(2), 247–255. 10.1038/s41386-019-0395-7

Cadoret, R. J., Troughton, E., O’Gorman, T. W., & Heywood, E. (1986). An adoption study of genetic and environmental factors in drug abuse. Archives of general psychiatry, 43(12), 1131–1136. 10.1001/archpsyc.1986.01800120017004

Canonica, T.& Zalachoras, I. (2022). Motivational disturbances in rodent models of neuropsychiatric disorders. Frontiers in Behavioral Neuroscience, 16(940672). 10.3389/fnbeh.2022.940672

Castro, N. C. F, Silva, I. S, Cartágenes, S. C., Fernandes, L. M. P., Ribera, P. C., Barros, M. A., Prediger, R. D., Fontes-Júnior, E. A., & Maia, C. S. F. (2022). Morphine Perinatal Exposure Induces Long-Lasting Negative Emotional States in Adult Offspring Rodents. Pharmaceutics, 14(1), 29. 10.3390/pharmaceutics14010029

Chiang, YC., Hung, TW., & Ho, IK. (2014). Development of sensitization to methamphetamine in offspring prenatally exposed to morphine, methadone and buprenorphine. Addiction Biology, 19(4), 676–686. 10.1111/adb.12055

Coyle, J. T. & Pert, C. B. (1976). Ontogenetic development of [3H]-naloxone binding in rat brain. Neuropharmacology, 15(1976), 555–560. 10.1016/0028-3908(76)90107-6

Desai, R. J., Hernandez-Diaz, S., Bateman, B. T., & Huybrechts, K. F. (2014). Increase in prescription opioid use during pregnancy among Medicaid-enrolled women. Obstetrics and gynecology, 123(5), 997–1002. 10.1097/AOG.0000000000000208

Gagin, R., Cohen, E., & Shavit, Y. (1996). Prenatal exposure to morphine alters analgesic responses and preference for sweet solutions in adult rats. Pharmacology, biochemistry, and behavior, 55(4), 629–634. 10.1016/s0091-3057(96)00278-x

Gardon, O., Faget, L., Chu Sin Chung, P., Matifas, A., Massotte, D., & Kieffer, B. L. (2014). Expression of mu opioid receptor in dorsal diencephalic conduction system: new insights for the medial habenula. Neuroscience, 277, 595–609. 10.1016/j.neuroscience.2014.07.053

Grecco, G. G., & Atwood, B. K. (2020). Prenatal Opioid Exposure Enhances Responsiveness to Future Drug Reward and Alters Sensitivity to Pain: A Review of Preclinical Models and Contributing Mechanisms. eNeuro, 7(6), ENEURO.0393-20.2020. 10.1523/ENEURO.0393-20.2020

Handelmann, G. E., & Quirion, R. (1983). Neonatal exposure to morphine increases mu opiate binding in the adult forebrain. European journal of pharmacology, 94(3-4), 357–358. 10.1016/0014-2999(83)90429-6

Harder, H. J., & Murphy, A. Z. (2019). Early life opioid exposure and potential long-term effects. Neurobiology of stress, 10, 100156. 10.1016/j.ynstr.2019.100156

Harder H. J., Searles C. T., Vogt M. E., Murphy A. Z. (2023). Perinatal opioid exposure leads to decreased social play in adolescent male and female rats: Potential role of oxytocin signaling in brain regions associated with social reward. Hormones and Behavior 153, 105384. 10.1016/j.yhbeh.2023.105384

Hayward, M. D., Schaich-Borg, A., Pintar, J. E., & Low, M. J. (2006). Differential involvement of endogenous opioids in sucrose consumption and food reinforcement. Pharmacology Biochemistry and Behavior, 85(3), 601–611. 10.1016/j.pbb.2006.10.015

Henderson-Redmond A. & Czachowski, C. (2014). Effects of systemic opioid receptor ligands on ethanol- and sucrose seeking and drinking in alcohol-preferring (P) and Long Evans rats. Psychopharmacology (Berl), 231(22), 4309–4321. 10.1007/s00213-014-3571-9

Heyser, C. J., Roberts, A. J., Schulteis, G., & Koob, G. F. (1999). Central administration of an opiate antagonist decreases oral ethanol self-administration in rats. Alcoholism, clinical and experimental research, 23(9), 1468–1476. 10.1111/j.1530-0277.1999.tb04669.x

Hyytiä, P., & Kiianmaa, K. (2001). Suppression of ethanol responding by centrally administered CTOP and naltrindole in AA and Wistar rats. Alcoholism, clinical and experimental research, 25(1), 25–33. 10.1111/j.1530-0277.2001.tb02123.x

Kocherlakota, P. (2014). Neonatal abstinence syndrome. Pediatrics, 134(2), e547–e561. 10.1542/peds.2013-3524

Konijnenberg, C., Sarfi, M., Melinder, A. 2016). Mother-child interaction and cognitive development in children prenatally exposed to methadone or buprenorphine. Early Human Development, 101, 91–97. 10.1016/j.earlhumdev.2016.08.013

Koob, G. F. (2020). Neurobiology of Opioid Addiction: Opponent Process, Hyperkatifeia and Negative Reinforcement. Biological Psychiatry, 87(1), 44–53. 10.1016/j.biopsych.2019.05.023

Kuhn, C. M., Windh, R. T., & Little, P. J. (1992). Effects of perinatal opiate addiction on neurochemical development of the brain. M.W. Miller (Ed.), Development of the Central Nervous System: Effects of Alcohol and Opiates, Wiley-Liss, New York, 341–361.

Lachenmeier, D.W. & Rehm, J. (2015). Comparative risk assessment of alcohol, tobacco, cannabis and other illicit drugs using the margin of exposure approach. Scientific Reports, 5(8126). 10.1038/srep08126

Larson, J. J., Graham, D. L., Singer, L. T., Beckwith, A. M., Terplan, M., Davis, J. M., Martinez, J., & Bada, H. S. (2019). Cognitive and Behavioral Impact on Children Exposed to Opioids During Pregnancy. Pediatrics, 144(2), e20190514. 10.1542/peds.2019-0514

Lê, A., & Shaham, Y. (2002). Neurobiology of relapse to alcohol in rats. Pharmacology & therapeutics, 94(1-2), 137–156. 10.1016/s0163-7258(02)00200-0

Lee, JS., Lee, SB., Kim, DW., Shin, N., Jeong, SJ., Yang, CH., Son, CG. (2021). Social isolation– related depression accelerates ethanol intake via microglia-derived neuroinflammation. Science Advances, 7 (45) 10.1126/sciadv.abj3400

McBride, W. J., Chernet, E., McKinzie, D. L., Lumeng, L., & Li, T. K. (1998). Quantitative autoradiography of mu-opioid receptors in the CNS of alcohol-naive alcohol-preferring P and -nonpreferring NP rats. Alcohol (Fayetteville, N.Y.), 16(4), 317–323. 10.1016/s0741-8329(98)00021-4

Mechling, A. E., Arefin, T., Lee, H. L., Bienert, T., Reisert, M., Ben Hamida, S., Darcq, E., Ehrlich, A., Gaveriaux-Ruff, C., Parent, M. J., Rosa-Neto, P., Hennig, J., von Elverfeldt, D., Kieffer, B. L., & Harsan, L. A. (2016). Deletion of the mu opioid receptor gene in mice reshapes the reward-aversion connectome. Proceedings of the National Academy of Sciences of the United States of America, 113(41), 11603–11608. 10.1073/pnas.1601640113

Morales, I., Rodríguez-Borillo, O., Font, L., & Pastor, R. (2020). Effects of naltrexone on alcohol, sucrose, and saccharin binge-like drinking in C57BL/6J mice: a study with a multiple bottle choice procedure. Behavioral Pharmacology, 31(2&3):256-271. 10.1097/FBP.0000000000000553

Nurmi, M., Kiianmaa, K., & Sinclair, J. D. (1999). Brain ethanol levels after voluntary ethanol drinking in AA and Wistar rats. Alcohol (Fayetteville, N.Y.), 19(2), 113–118. 10.1016/s0741-8329(99)00022-1

Palamar, J. J., Le, A., & Mateu-Gelabert, P. (2018). Not just heroin: Extensive polysubstance use among US high school seniors who currently use heroin. Drug and alcohol dependence, 188, 377–384. 10.1016/j.drugalcdep.2018.05.001

Patrick, S. W., Davis, M. M., Lehman, C. U., & Cooper, W. O. (2015). Increasing incidence and geographic distribution of neonatal abstinence syndrome: United States 2009 to 2012. Journal of perinatology : official journal of the California Perinatal Association, 35(8), 667. 10.1038/jp.2015.63

Patrick, S. W., Faherty, L. J., Dick, A. W., Scott, T. A., Dudley, J., & Stein, B. D. (2019). Association Among County-Level Economic Factors, Clinician Supply, Metropolitan or Rural Location, and Neonatal Abstinence Syndrome. JAMA, 321(4), 385–393. 10.1001/jama.2018.20851

Paxinos G. & Watson C. (1997). The rat brain in stereotaxic coordinates. New York: Academic.

Peris, J., Zharikova, A., Li, Z., Lingis, M., MacNeill, M., Wu, M. T., & Rowland, N. E. (2006). Brain ethanol levels in rats after voluntary ethanol consumption using a sweetened gelatin vehicle. Pharmacology, biochemistry, and behavior, 85(3), 562–568. 10.1016/j.pbb.2006.10.010

Ramsey, N. F., Niesink, R. J., & Van Ree, J. M. (1993). Prenatal exposure to morphine enhances cocaine and heroin self-administration in drug-naive rats. Drug and alcohol dependence, 33(1), 41–51. 10.1016/0376-8716(93)90032-l

Roger-Sánchez C., Aguilar M. A., Rodríguez-Arias M., Aragon C. M., & Miñarro J. (2012). Age- and sex-related differences in the acquisition and reinstatement of ethanol CPP in mice. Neurotoxicology and Teratology, 34(1),108–15. 10.1016/j.ntt.2011.07.011

Russell, B. S., Trudeau, J. J., & Leland, A. J. (2015). Social Influence on Adolescent Polysubstance Use: The Escalation to Opioid Use. Substance use & misuse, 50(10), 1325– 1331. 10.3109/10826084.2015.1013128

Schindler, C. J., Slamberová, R., Rimanóczy, A., Hnactzuk, O. C., Riley, M. A., & Vathy, I. (2004). Field-specific changes in hippocampal opioid mRNA, peptides, and receptors due to prenatal morphine exposure in adult male rats. Neuroscience, 126(2), 355–364. 10.1016/j.neuroscience.2004.03.040

Shen, Y. L., Chen, S. T., Chan, T. Y., Hung, T. W., Tao, P. L., Liao, R. M., Chan, M. H., & Chen, H. H. (2016). Delayed extinction and stronger drug-primed reinstatement of methamphetamine seeking in rats prenatally exposed to morphine. Neurobiology of learning and memory, 128, 56–64. 10.1016/j.nlm.2015.12.002

Stromberg, M. F., Meister, S., Volpicelli, J. R., & Ulm, R. R. (1997). Morphine enhances selection of both sucrose and ethanol in a two-bottle test. Alcohol, 14(1), 55–62. 10.1016/s0741-8329(96)00107-3

Substance Abuse and Mental Health Services Administration (SAMHSA). 2015 National Survey on Drug Use and Health (NSDUH). Table 2.19B: Alcohol Use in Lifetime, Past Year, and Past Month, by Detailed Age Category:Percentages, 2014 and 2015. Rockville, MD: SAMHSA, 2016. Available at: http://www.samhsa.gov/data/sites/default/files/NSDUH-DetTabs-2015/NSDUH-DetTabs-2015/NSDUH-DetTabs-2015.htm#tab2-19b

Torabi, M., Pooriamehr, A., Bigdeli, I., & Miladi-Gorji, H. (2017). Maternal swimming exercise during pregnancy attenuates anxiety/depressive-like behaviors and voluntary morphine consumption in the pubertal male and female rat offspring born from morphine dependent mothers. Neuroscience letters, 659, 110–114. 10.1016/j.neulet.2017.08.074

Vassoler, F. M., Wright, S. J., & Byrnes, E. M. (2016). Exposure to opiates in female adolescents alters mu opiate receptor expression and increases the rewarding effects of morphine in future offspring. Neuropharmacology, 103, 112–121. 10.1016/j.neuropharm.2015.11.026

Vathy, I., Slamberová, R., & Liu, X. (2007). Foster mother care but not prenatal morphine exposure enhances cocaine self-administration in young adult male and female rats. Developmental psychobiology, 49(5), 463–473. 10.1002/dev.20240

Vathy, I., Slamberová, R., Rimanóczy, A., Riley, M. A., & Bar, N. (2003). Autoradiographic evidence that prenatal morphine exposure sex-dependently alters mu-opioid receptor densities in brain regions that are involved in the control of drug abuse and other motivated behaviors. Progress in neuro-psychopharmacology & biological psychiatry, 27(3), 381–393. 10.1016/S0278-5846(02)00355-X

Watson R. E., Wiegand S. J., Clough R. W., Hoffman G. E. (1986). Use of cryoprotectant to maintain longterm peptide immunoreactivity and tissue morphology. Peptides, 7(1), 155–159. 10.1016/0196-9781(86)90076-8

Welle-Strand, G. K., Skurtveit, S., Jansson, L. M., Bakstad, B., Bjarkø, L., & Ravndal, E. (2013). Breastfeeding reduces the need for withdrawal treatment in opioid-exposed infants. Acta paediatrica (Oslo, Norway : 1992), 102(11), 1060–1066. 10.1111/apa.12378

Winkelman, T., Villapiano, N., Kozhimannil, K. B., Davis, M. M., & Patrick, S. W. (2018). Incidence and Costs of Neonatal Abstinence Syndrome Among Infants With Medicaid: 2004-2014. Pediatrics, 141(4), e20173520. 10.1542/peds.2017-3520

Wu, L. Y., Chen, J. F., Tao, P. L., & Huang, E. Y. (2009). Attenuation by dextromethorphan on the higher liability to morphine-induced reward, caused by prenatal exposure of morphine in rat offspring. Journal of biomedical science, 16(1), 106. 10.1186/1423-0127-16-106

Xu, P., Wang, K., Lu, C., Dong, L., Chen, Y., Wang, Q., Shi, Z., Yang, Y., Chen, S., z7 Liu, X. (2017). Effects of the chronic restraint stress induced depression on reward-related learning in rats. Behavioural brain research, 321,185–192. 10.1016/j.bbr.2016.12.045

