## Supplementary figures and images for "Perigestational Opioid Exposure Alters Alcohol-Driven Reward Behaviors in Adolescent Rats"

### Supplemental Figures

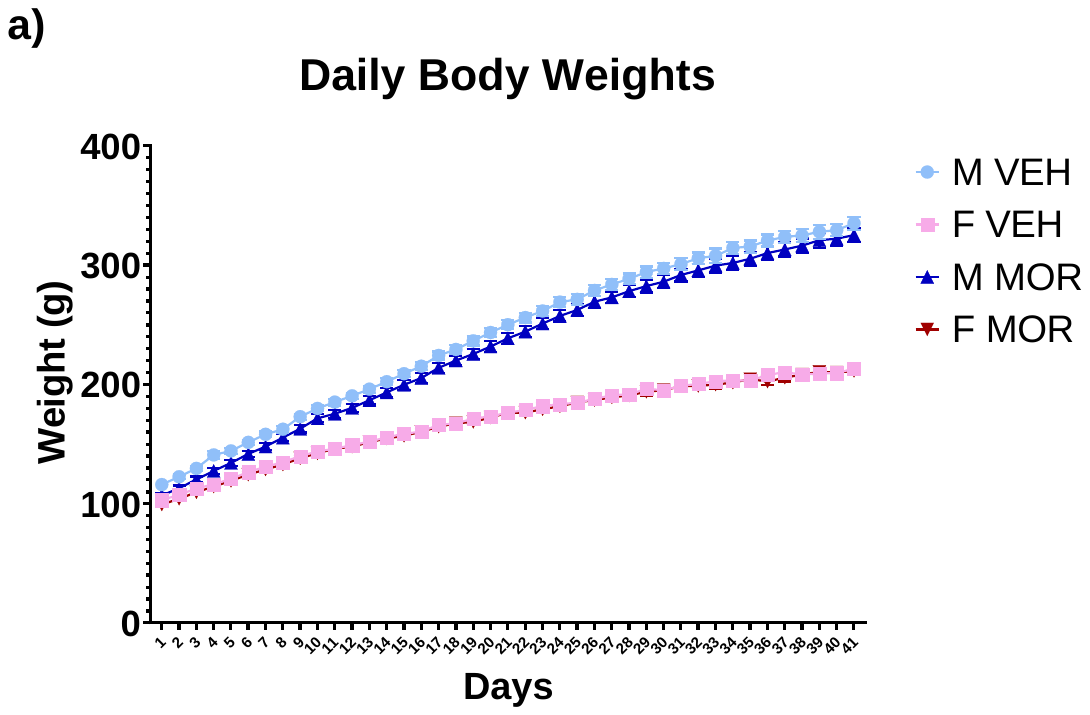

Supplemental Figure 1.

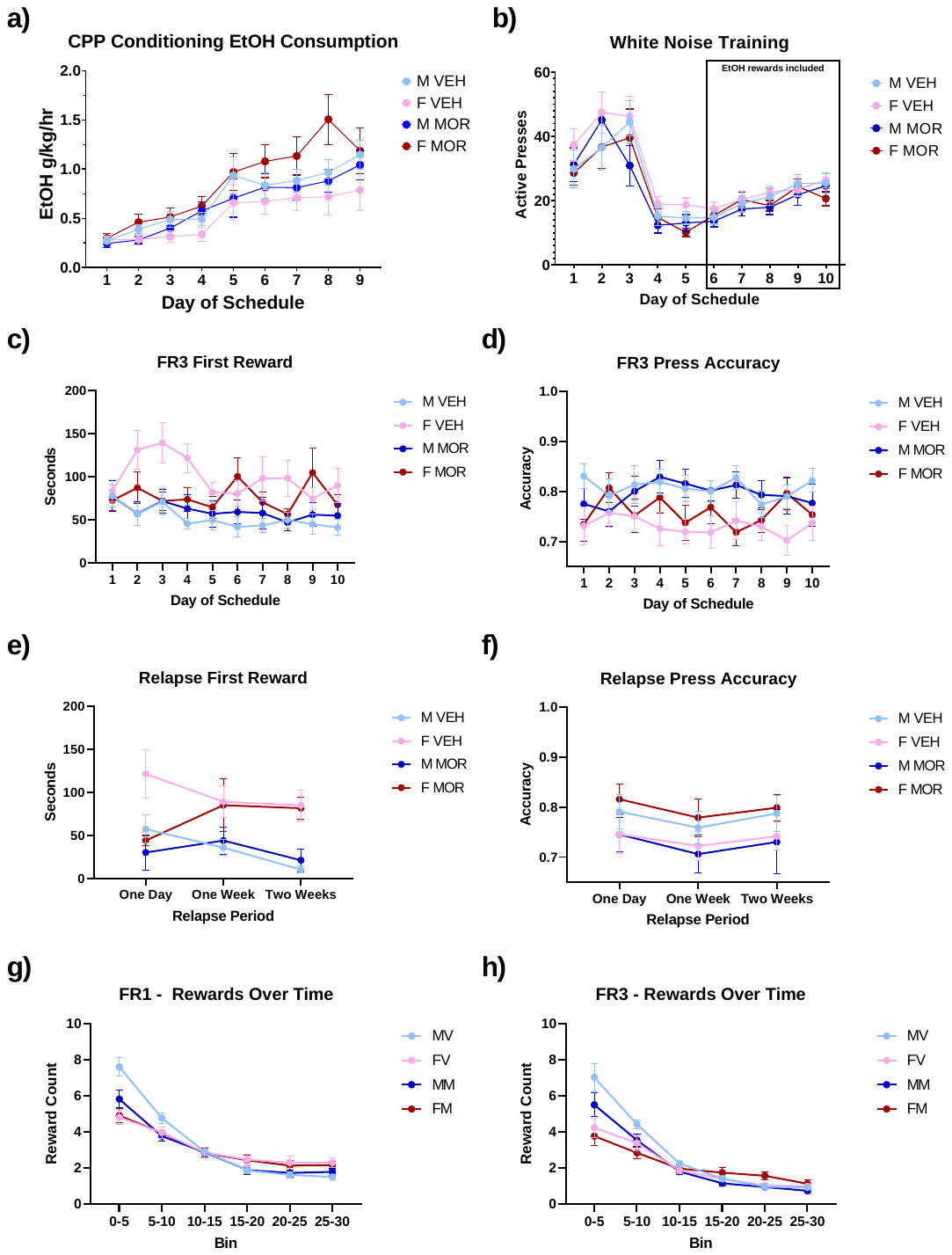

Supplemental Figure 2.

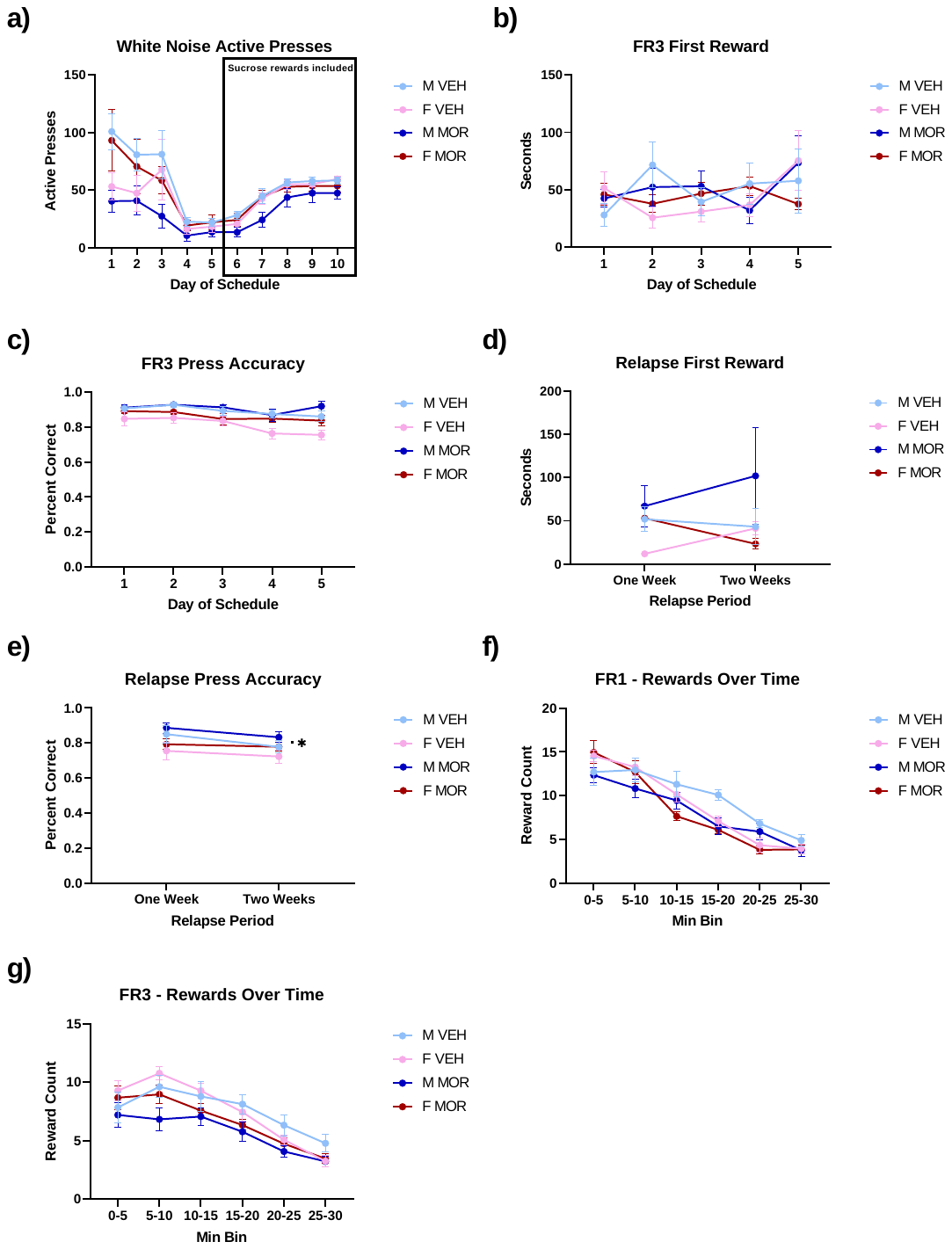

Supplemental Figure 3.
